# Kidney pathology alters cell-clustering in single cell RNA sequencing

**DOI:** 10.1101/2021.10.08.463030

**Authors:** Lijun Ma, Mariana Murea, Young A Choi, Ashok K. Hemal, Alexei V. Mikhailov, James A. Snipes, Jeff W. Chou, Robert C. Langefeld, Wei Cui, Lance D. Miller, Gregory A. Hawkins, Nicholette D. Palmer, Carl D. Langefeld, Barry I. Freedman

**Author notes:** Corresponding authors: Lijun Ma, MD, PhD or Barry I. Freedman, MD, Department of Internal Medicine, Section on Nephrology, Wake Forest School of Medicine, Medical Center Boulevard, Winston Salem, NC 27157 USA, or.

## Abstract

Single cell RNA sequencing (scRNA-Seq) is useful to classify cell-specific gene expression profiles in kidney tissue. As viable cells are required, we report an optimized cell dissociation methodology and the necessity of screening tissue histology prior to scRNA-Seq. We demonstrate that glomerular injury can selectively reduce the appearance of groups of cells during analysis of cell clustering and we confirmed reductions in cell-specific markers among injured cells on kidney sections with fluorescence microscopy. Interpretation of scRNA-Seq results may be refined based upon these considerations.

## Introduction

Due to the heterogeneity of cell types in the kidney, researchers have proposed and performed microdissection-based expression quantitative trait loci (eQTL) analyses on human kidney cell fractions[1]. More recently, single-cell RNA sequencing (scRNA-Seq) has been recognized as a cutting-edge technique in the classification of cell-specific gene expression profiles, particularly important to determine cell-specific mechanisms underlying kidney disease.

Kidney biopsies are required to perform cell-specific kidney tissue eQTL analysis or differential gene expression studies. It is crucial to maintain cell viability for scRNA-Seq; however, many cells do not survive tissue processing. This is likely the case in patients with advanced chronic glomerular diseases, including focal segmental glomerulosclerosis (FSGS), and prior to transplant of deceased donor kidneys with prolonged cold ischemia. Based on the need for viable cell populations for scRNA-Seq, these concerns may support the assessment of kidney histology from nephrectomy or biopsy specimens prior to analysis.

## Methods

### Collection of nephrectomy specimens

We randomly selected four patients in our repository who underwent nephrectomy for renal cell carcinoma between 8/23/2013 and 9/26/2013 to verify the quality of enriched glomerular cells after long-term storage in liquid nitrogen (LN2). Patient age, sex, diabetes status, and post-surgical complications were recorded. Preoperative serum creatinine and estimated glomerular filtration rate (eGFR) were obtained. All patients had a preoperative CKD-EPI eGFR >60 ml/min/1.73 m^2^ Basic demographic data are displayed in Supplementary Table 1. The Institutional Review Board at the Wake Forest School of Medicine approved the protocol and all patients provided written informed consent.

### Specimen processing and sc-RNA-Seq

After nephrectomy, cortical tissue was immediately dissected in the operating room to obtain non-diseased tissue. Macroscopically healthy tissue was immediately washed with sterile Hank’s balanced salt solution (HBSS; Lonza, Walkersville, MD), and portions were: a) placed in a biopsy cassette (Fisher HealthCare, Houston, TX) and immersed in 4% paraformaldehyde for preparation of formalin-fixed paraffin-embedded (FFPE) kidney tissue blocks for subsequent histologic examination (Supplementary Materials), b) rapidly frozen in optimal cutting temperature gel (Sukaru FineTek Inc., Torrance, CA) to create cryosections for immunofluorescence studies (Supplementary Materials), and c) kept in Dulbecco’s Modified Eagle Medium (DMEM; Invitrogen, Grand Island, NY) media with 10% fetal bovine serum (FBS) on a surface of ice or 4°C refrigeration for up to 36 hours (mimicking the maximum time for transplantation after recovery of donor kidneys) prior to the next step in glomerular enrichment.

For glomerular cell enrichment, kidney tissues were washed on ice in HBSS (Lonza, Walkersville, MD) after removal from DMEM. Cortical tissue was cut into 1 mm^3^ fragments and digested with 1 mg/ml collagenase type II at 37° C in DMEM (Invitrogen, Grand Island, NY) for 30 minutes. The suspension was passed through a 250 μm sieve (Sigma, St. Louis, MO) with minimum pressure. The flow through was subsequently passed through a 100 μm cell strainer (BD BioSciences). Glomeruli were collected on the surface of the strainer. After three washes with HBSS, glomeruli were suspended in 1 ml DMEM containing 0.5% BSA in a 1.5 ml centrifuge tube, and incubated at room temperature for 15 minutes with gentle agitation. Enrichment of glomeruli were assessed via light microscopy using a drop of the suspension (Supplementary Figure 1). The glomerular suspension was digested with 0.5 mg/ml of collagenase (Invitrogen), 0.5 mg/ml dispase II (ZenBio, Research Triangle Park, NC), and 0.075% trypsin in DMEM at 37° C for 20 minutes with gentle rotation. Detached cells were treated with DMEM containing 10% FBS to neutralize proteases and centrifuged at 300 x g for 3 minutes. The cell pellet was re-suspended and cultured in Endothelial Cell Growth Media (EGM2)-MV (Lonza, Walkersville, MD) for a two-day recovery before washing in HBSS and trypsinization. Cells were viably frozen in DMEM with 20% heat-inactivated FBS and 10% Hybri-Max Dimethyl Sulfoxide (DMSO; Sigma-Aldrich): by cooling in isopropanol at −1°C per minute at −80°C overnight, and subsequently stored under LN2 vapor. In preparation for scRNA-seq, cells were thawed and washed according to the protocol for human peripheral blood mononuclear cells (PBMCs; 10× Genomics). All scRNA-seq procedures were performed by the Cancer Genomics Shared Resource (CGSR) of Wake Forest School of Medicine. Viable cells (mean 85.35% ± 2.3%, n=4) in suspensions averaging 1205 ± 130 cell/μl were loaded into wells of a 10× Chromium single cell capture chip targeting a cell recovery rate of 2000-4000 cells. Single-cell gel beads in emulsion (GEMs) were created on a Chromium Single Cell Controller and scRNA-seq libraries were prepared using the Chromium Single Cell 3’ Library and Gel Bead kit according to the manufacturer’s protocol (10× Genomics). Sequencing libraries were loaded on an Illumina NovaSeq 6000 with High Output 150 cycle kit (Illumina) for paired-end sequencing using the following read length: 26 bp Read1, 8 bp i7 Index, 0 bp i5 Index, and 98 bp Read2). Technical procedures are highlighted in Supplementary Figure 1.

### Processing of scRNA-Seq data

Cell Ranger Single Cell Software Suite v.3.0.1 was used to perform sample de-multiplexing, alignment, filtering, and UMI (*i.e.*, unique molecular identifier) counting (https://support.10xgenomics.com/single-cell-gene-expression/software/pipelines/latest/algorithms/overview). The data for each respective subpopulation were aggregated for direct comparison of single cell transcriptomes. A total of 11,259 single cells from the 4 samples were captured, with the number of cells recovered per channel ranging from 2486 to 3546. The mean reads per cell varied from 63,761 and 88,599 with median Unique Molecular Indexes of 3,429 to 5,257 per cell. The ScRNA seq data analysis pipeline is described in Supplementary Materials.

## Results

### Histologic findings on hematoxylin and eosin-stained FFPE kidney sections

Figure 1 displays normal glomerular and tubular structures in patients A, B and D. Patient A had long-standing diabetes. The biopsy from Patient C (Figure 1) showed dilated afferent and efferent arterioles, as well as dilated capillaries in the glomerular tuft and peritubular regions packed with red blood cells. Mild to moderate diffuse acute tubular injury was present. Potential causes of this histology include venous outflow obstruction due to compression of the renal veins by tumor, venous thrombosis related to paraneoplastic effects, systemic disease, or thrombotic microangiopathy. Although diagnostic features of thrombotic microangiopathy were absent, the patient subsequently developed deep vein thrombosis after the nephrectomy.

**Figure 1.**
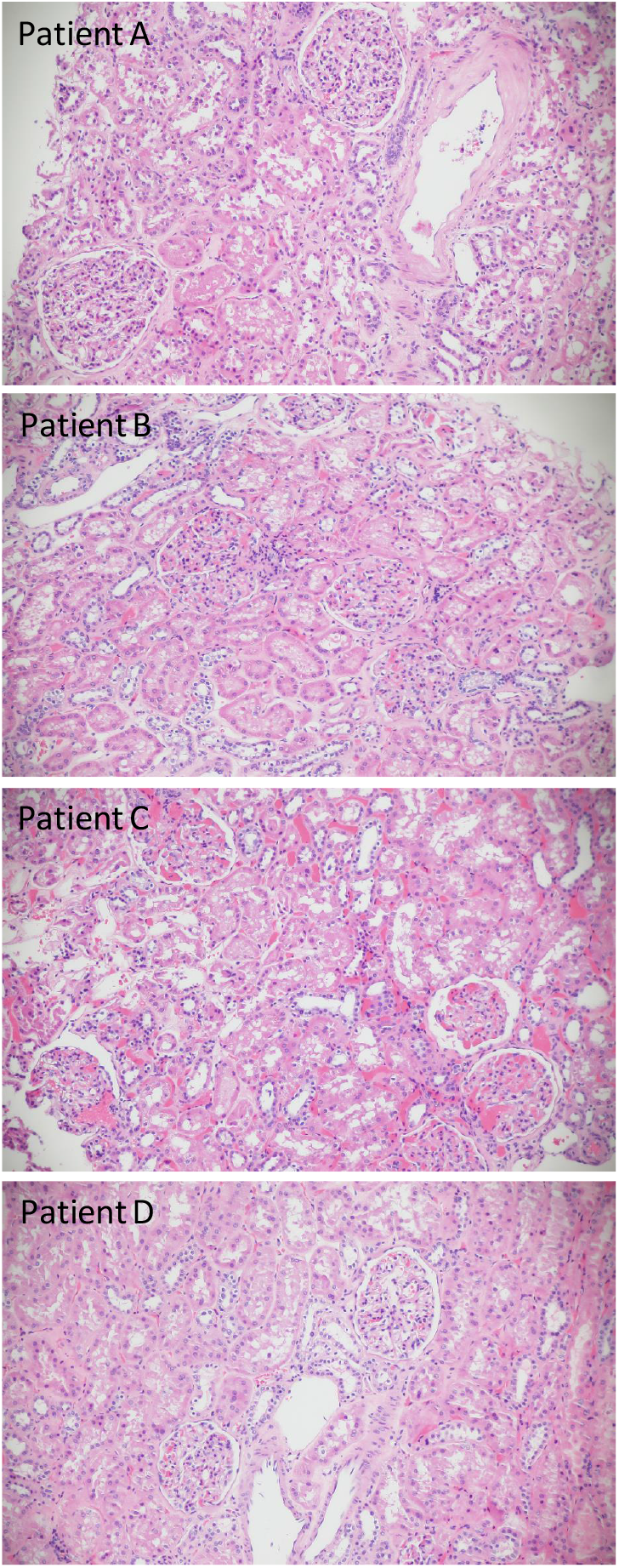
Hematoxylin and Eosin staining of FFPE kidney sections. Patients A, B and D display normal glomeruli and tubules. Patient C has congested arterioles and peritubular capillaries, with acute tubular injury (All x20).

### Kidney cell types, grouped by scRNA Seq features

Figure 2a displays the major kidney cell types in the study participants, grouped by cell-specific markers. The top 5 transcripts for each cell cluster are summarized based on FDR p-values in Supplementary Table 2. Two-part hurdle FDR was used to account for cells from the same individual and validate the significance of original FDR. Podocytes were classified by a number of known markers, with *NPHS2* (podocin) as the top marker with FDR p-value=6×10^-265^ (log_2_[fold-change]=9.39). Glomerular endothelial cells were readily clustered by known markers, with *PECAM1* (CD31) ranking highest for its FDR p-value=9×10^-151^ (log_2_[fold-change]=9.76). Another cluster of endothelial cells was detected, featured by top marker *VCAM1* with FDR p-value=6.0×10^-38^ (log_2_[fold-change]=2.23). *VCAM1* encodes a cell surface sialoglycoprotein expressed by cytokine-activated endothelial cells. Mesangial cells were characterized by a series of markers, where *HHIP* (hedgehog interacting protein) was at the top with an FDR p-value=6×10^-114^ (log_2_[fold-change]=3.35). *HHIP* is a novel intracellular mesangial cell marker, which interacts with TGFβ1 and plays a role in mesangial expansion with diabetic kidney disease[2].

**Figure 2.**
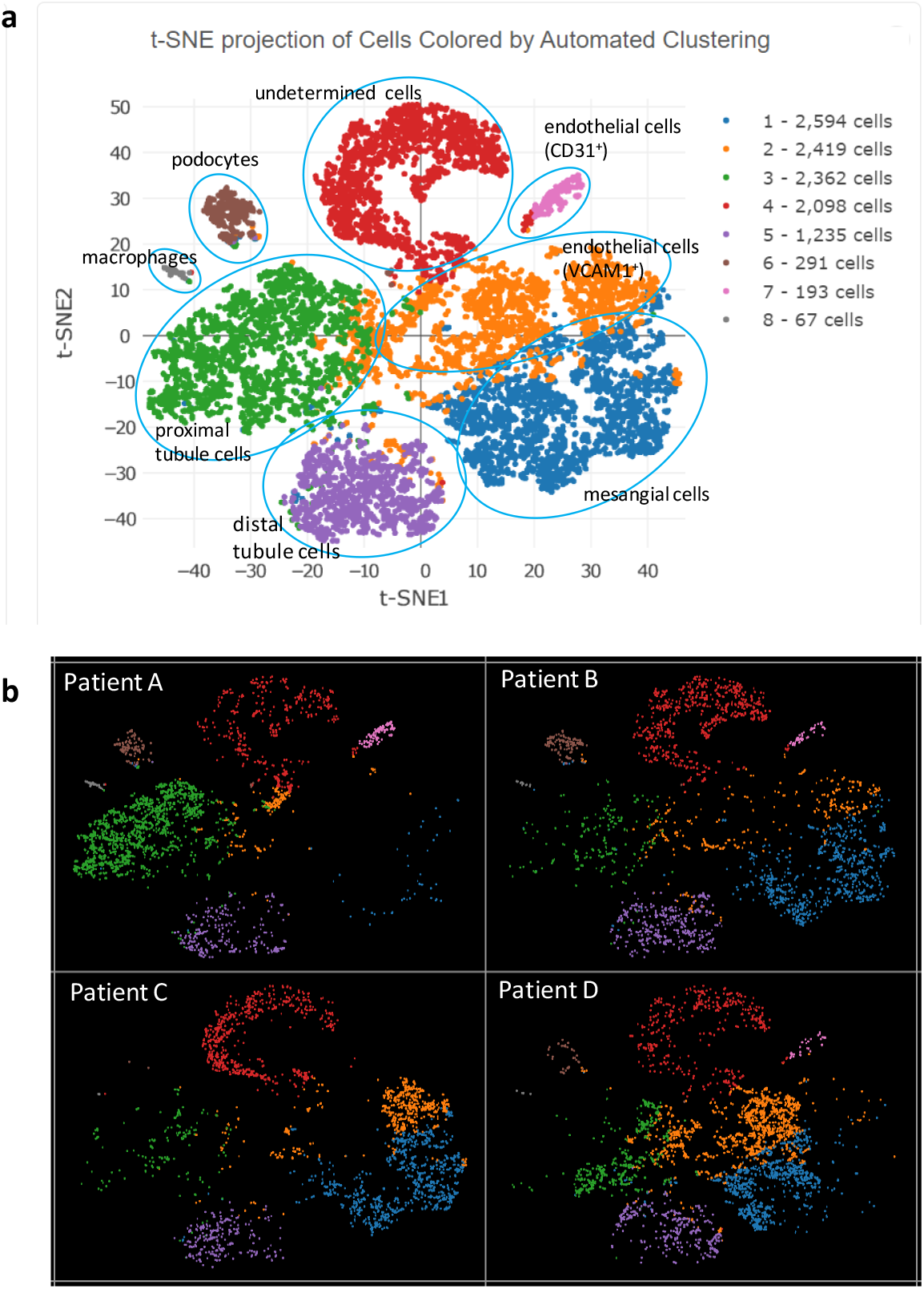
Feature cluster plot of enriched glomerular cells. a) Aggregation of cells from all 4 participants; b) cell clustering for individual participants.

A key observation from review of individual feature plots was that podocin-positive cells and CD31-positive glomerular endothelial cells were absent from Patient C (Figure 2b). We suspect that glomerular endothelial cell injury resulting from circulatory changes led to podocyte dysfunction in this individual. Patient C had well clustered proximal tubule and distal tubule cells, as seen in the other three patients in the aggregated feature plot (Figure 2 a, b) based on their corresponding cell markers, *i.e., ANPEP* (Aminopeptidase N, CD13) as top proximal tubule marker[3] (FDR p-value=5×10^-123^, log_2_[fold-change]=3.44) and *RAB25* as top distal tubule and collecting duct marker[4] (FDR p-value=2×10^-299^, log_2_[fold-change]=6.31). For the aggregated feature plot from all four patients, the macrophage cluster was readily detected by top markers *C1QA, C1QB*, and *C1QC* with FDR p-values=5×10^-157^, 3×10^-154^, and 2×10^-139^, respectively; log_2_[fold-change] 11.80 for all. The final group of cells were classified as “undetermined” (Figure 2a, Supplementary Table 2), featured by a group of overexpressed mitochondrion-encoded genes (Supplementary Figure 2). Lactate dehydrogenase *LDHA* expression levels in these cells were lower than among other cell clusters, indicating metabolic inactivity (Supplementary Figure 3). Therefore, some of these cells may have been dying (85% of cells in this pool were initially viable) or be latent glomerular progenitor cells that could develop into future glomerular cells[5] or podocytes[6]. We did not remove these cells to avoid the potential for bias, instead marking them “undetermined.”

### Verification of impacted cell clusters by immunofluorescence on kidney cryo-sections

To assess the discrepancies of cell grouping patterns across the four kidney biopsy samples in the scRNA-Seq feature plot (lack of podocyte and glomerular endothelial cell clusters in patient C), the immunofluorescence staining of podocin, CD31 and VCAM1 was examined using cryo-sections from the same four patients after HE staining of FFPE-sections and scRNA-Seq of glomerular cell-enriched cell suspensions. Podocin and CD31 signals were markedly lower in Patient C than in other three patients; however, they displayed typical glomerular podocyte and endothelial cell patterns, respectively, as in Patients A, B, and D (Supplementary Figures 4 & 5). Across all four patients, VCAM1 appeared to display Bowman’s capsule and peritubular endothelial cell patterns (Supplementary Figure 6), consistent with prior reports that VCAM-1 is present on parietal epithelial cells lining Bowman’s capsule[7] and peritubular endothelial cells[8].

## Discussion

Kidney specimens comprising enriched glomerular cells, cryo-tissue blocks, and FFPE tissue blocks collected nearly eight years prior were selected to assess effects of LN2 preservation time on patterns of scRNA-Seq cell clustering. The importance of scRNA-Seq in human kidney tissue is critical to detect disease mechanisms. Glomerular cells occupy a small fraction of total kidney volume; as such, scRNA-Seq in kidney compartments (cortex or interstitium) or cell collections that are not enriched for glomerular cells may result in failure to detect podocytes and other glomerular cells due to dilution with massive volumes of tubule cells during analysis of clustering data [9]. Microdissection requires trained personnel to rapidly complete the process; however, cell dissociation is still required[1]. Fluorescence-activated cell sorting (FACS)[10] prior to scRNA-Seq is feasible for small numbers of samples. To perform scRNA-Seq in larger studies (*e.g*., >50 individuals), FACS would be time-consuming, expensive, and difficult to complete in a narrow time frame. This could lead to batch effects.

Previous reports of scRNA-Seq in glomerular-enriched cells were performed shortly after FACS sorting, without time for cell recovery. In contrast, our protocol provided cells two additional days in recovery media. This led to high rates of cell survival after 8 years of storage in LN2. The modified cell dissociation method we applied avoided FACS[10], and obtained similar cell clusters after scRNA-Seq as those with FACS. Although proximal and distal tubule cell numbers were higher in the clustering feature plot, this did not appear to affect the quality of glomerular cell clustering.

Histology from Patient C showed markedly dilated arterioles and capillaries packed with red blood cells and acute tubular injury. Abnormal glomerular blood flow can lead to endothelial injury and injured podocytes[11]. To assess the viability of all glomerular-enriched cells and to assess the validity of cells in the scRNA-Seq analysis, we performed cell dissociation of enriched glomerular cell mixtures from all four patients. The overall scRNA-Seq cell clustering plot revealed the major glomerular cell types with reasonable mixtures of proximal tubule and distal tubule cells. However, podocin-positive and CD31-positive cells, representing podocytes and glomerular endothelial cells, respectively, were absent from the scRNA-Seq cell cluster breakdown plot from Patient C. Immunofluorescence on kidney cryosections in the combined four patients showed weaker podocin and CD31 signals in Patient C, suggesting compromised podocyte and glomerular endothelial cell function. We suspect altered glomerular perfusion resulted in podocyte dysfunction and stressed glomerular endothelial cells. As a result, podocytes may have been less likely to survive enzyme (protease) treatment during cell dissociation. Hence, they would be more difficult to preserve, resulting in loss of cells with positive glomerular endothelial cell and podocyte markers on the scRNA-Seq cell cluster feature plot.

This study demonstrates the need to consider kidney histology when performing scRNA-Seq. Histologic analysis can screen samples for kidney diseases that may reduce the viability of select cells after cell dissociation and long-term storage in LN2. Specific glomerular cell marker signals were verified with immunofluorescence on kidney cryo-sections; this more likely captured the native form of immunogens.

A limitation of this study is that only four kidney biopsies were selected for exploratory analyses. One displayed abnormal arteriolar and microcirculatory compartments, with associated tubular epithelial damage. The scRNA-Seq feature plot was consistent with glomerular ischemic injury. As scRNA-Seq technology is increasingly popular, the kidney community needs to be aware of potential confounders. In addition, we highlight our modified cell dissociation protocol that avoided the need to transport cryo-preserved samples to central laboratories[12] and bypasses FACS to simplify cell processing for larger scale clinical biopsy-based cell-specific glomerular gene expression analyses. This approach preserves cell viability by adding a recovery stage between cell dissociation and a single-time cryo-preservation of glomerular-enriched cells in LN2, allowing future scRNA-Seq analysis to be uniformly conducted in batches with short time windows to minimize technical variation. Single nuclear RNA sequencing may avoid the requirement to assess healthy viable cells; however, mature cytosolic transcript isoforms and mitochondrial encoded transcripts will not be detected by this method. Hence, for kidney diseases resulting from mitochondrial dysfunction, such as Mitochondrial Encephalopathy, Lactic Acidosis, and Stroke-like episodes (MELAS) syndrome and recently recognized *APOL1*-associated kidney disease, scRNA-Seq may be preferable.

We conclude histologic screening of kidney biopsy samples should be performed before scRNA-Seq analysis. Retrieving a relatively viable kidney cell pool is critical to using cell-specific gene expression profiles from scRNA-Seq to investigate mechanisms underlying kidney disease.

## Statement of Ethics

These experiments were performed in accordance with the Institutional Review Board (IRB) at Wake Forest School of Medicine. All criteria set by the IRB were met.

## Author Contributions

L.M., C.D.L., and B.I.F. designed the study. L.M. and B.I.F. drafted and revised the manuscript. M.M. and A.K.H. recruited the patients. A.V.M. and M.M. reviewed histology results and critically interpreted the data and review the manuscript. Y.A.C. and J.A.S performed cell biology and immunofluorescence experiments. W.C. performed the scRNA seq experiment. J.W.C. and R.C.L. analyzed the scRNA seq data. C.D.L. supervised the data analysis. L.C.D. and G.A.H. reviewed the analysis. All authors reviewed the paper and approved the final version of the manuscript.

## Data Availability

The scRNA-seq raw data are openly available at the NCBI sequence read archive (https://www.ncbi.nlm.nih.gov/sra/PRJNA781289)

## Supplementary Materials

### Supplementary Methods

#### Histological examination and immunofluorescence imaging

Prior to thawing the glomerular-enriched cells frozen in liquid nitrogen vapor for single cell RNA sequencing (scRNA Seq) analysis, formalin-fixed paraffin-embed (FFPE) kidney tissue blocks were sliced into 3 μm thickness sections for staining with hematoxylin and eosin (HE); digital images were obtained on a Zeiss Axioplan 2 microscope. Immunofluorescence of podocin (Santa Cruz Biotechnology), CD31 (BD Biosciences), and VCAM1 (Invitrogen) was performed on kidney cryo-sections using established protocols[1] from patients who underwent HE staining of FFPE sections for histologic examination. Information for primary antibodies is summarized in Supplementary Table 3. Secondary antibodies (donkey anti-goat Alexa Fluor 594, and goat anti-mouse Alexa Fluor 488, Jackson ImmunoResearch Laboratories) were used to display fluorescent signals (1:100 dilution). Immunofluorescence imaging was performed on an Olympus IX71 fluorescence microscope (Olympus Scientific Solutions Americas Corp., Waltham, Massachusetts).

#### Pipeline for ScRNA Seq data analysis

To reduce the gene expression matrix to its key features, Cell Ranger employs Principal Components Analysis (PCA) to reduce the dimensionality of the dataset from (cells x genes) to (cells x N), where N is the number of principal components. As detailed in a technical review from 10x Genomics (https://support.10xgenomics.com/single-cell-gene-expression/software/pipelines/latest/algorithms/overview), the analysis pipeline uses a Python implementation of the IRLBA algorithm[2], designed to reduce memory consumption. For T-distributed Stochastic Neighbor Embedding (tSNE) projection and clustering analysis, the first 30 principal components were used, determined by the visualized PCA Elbow plot in the Seurat package.

To identify genes whose expression is specific to each cluster, the two-part hurdle mixed model, with individual (person) as a random effect, as implemented in MAST[3] was computed[4]. The two-part hurdle model jointly tests two hypotheses: 1) proportion of cells expressing the transcript, and 2) given gene is expressed, the magnitude of expression. In this generalized linear mixed model, the random effect accounts for the within individual correlation in gene expression and the two-part hurdle accounts for zero inflation. Because the number of individuals is modest (i.e., four individuals), some applications of the two-part hurdle model will not converge and the pseudobulk analysis as implemented in Cell Ranger is reported. Specifically, Cell Ranger tested whether the mean expression (for every transcript and every cluster) in cluster *k* differed from the mean expression across all other cells. For cluster *k*, the mean expression of a featured transcript is calculated as the total number of UMIs from that transcript in cluster *k* divided by the sum of the size factors for cells in cluster *k*. The size factor for each cell is the total UMI counts in that cell divided by the median UMI count per cell (across all cells). The mean expression outside of cluster *k* is calculated in the same way. The log_2_ fold-change of expression in cluster *k* relative to other clusters is defined as the log_2_ ratio of mean expression within cluster *k* and outside of cluster *k*. When computing the log_2_ fold-change, a pseudocount of 1 is added to both the numerator and denominator of the mean expression.

To test for differences in mean expression between groups of cells, Cell Ranger uses the exact negative binomial test proposed by the sSeq method[5]. When the counts become large, Cell Ranger switches to the fast asymptotic negative binomial test used in edgeR[6].

The feature plot function was used to highlight cell clusters expressing specific transcripts. A false discovery rate (FDR) p-value and fold change were used to estimate the differential expression of transcripts grouped in the specific cluster vs. all other cells.

---

**Supplementary Table 1.** Demographic data in study participants at nephrectomy

| **Patient** | **Age**<br>(years) | **Sex** | **BMI<sup>1</sup>**<br>(kg/m<sup>2</sup>) | **eGFR<sup>2</sup>**<br>(ml/min/1.73m<sup>2</sup>) | **Screat<sup>3</sup>**<br>(mg/dL) | **Diabetes** |
| --- | --- | --- | --- | --- | --- | --- |
| A | 64 | F | 34.1 | >60 | 0.57 | yes |
| B | 42 | M | 30.2 | >60 | 0.89 | no |
| C | 49 | M | 29.1 | >60 | 1.07 | no |
| D | 39 | M | 24.0 | >60 | 0.93 | no |

<sup>1</sup> body mass index
<sup>2</sup> estimated glomerular filtration rate based on CKD-EPI Equation (2021)
<sup>3</sup> serum creatinine concentration

**Supplementary Table 2.**
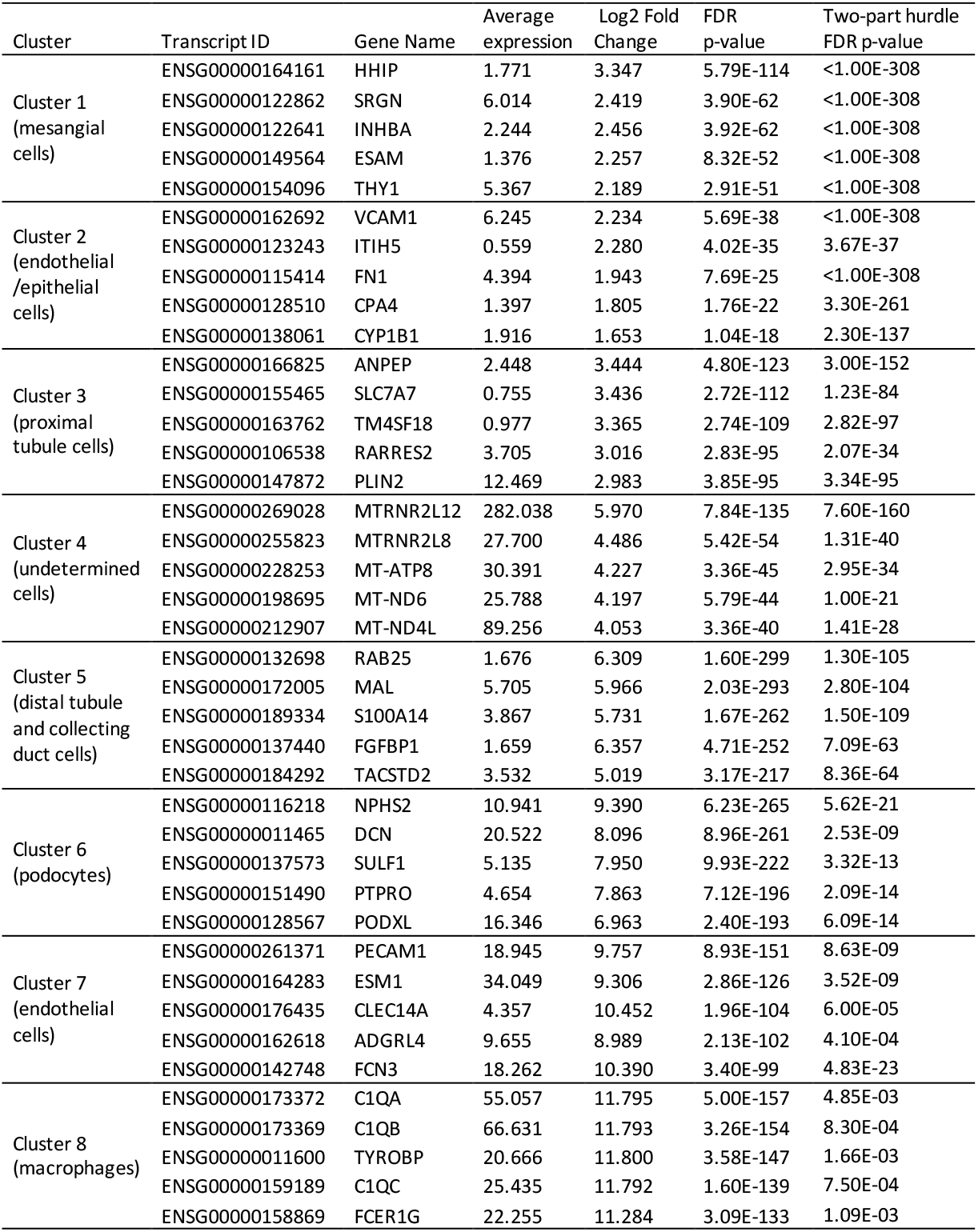
Top 5 transcripts in each cell cluster

**Supplementary Table 3.** Primary antibodies used in immunofluorescence microscopy

| Antibody | Vendor | Catlog # | Host species | Dilution |
| --- | --- | --- | --- | --- |
| Podocin | Santa Cruz | sc22294 | goat | 1:50 |
| CD31(PECAM1) | BD BioSciences | 550389 | mouse | 1:25 |
| VCAM1 | Invitrogen | 14-1069-82 | mouse | 1:100 |

**Supplementary Figure 1.**
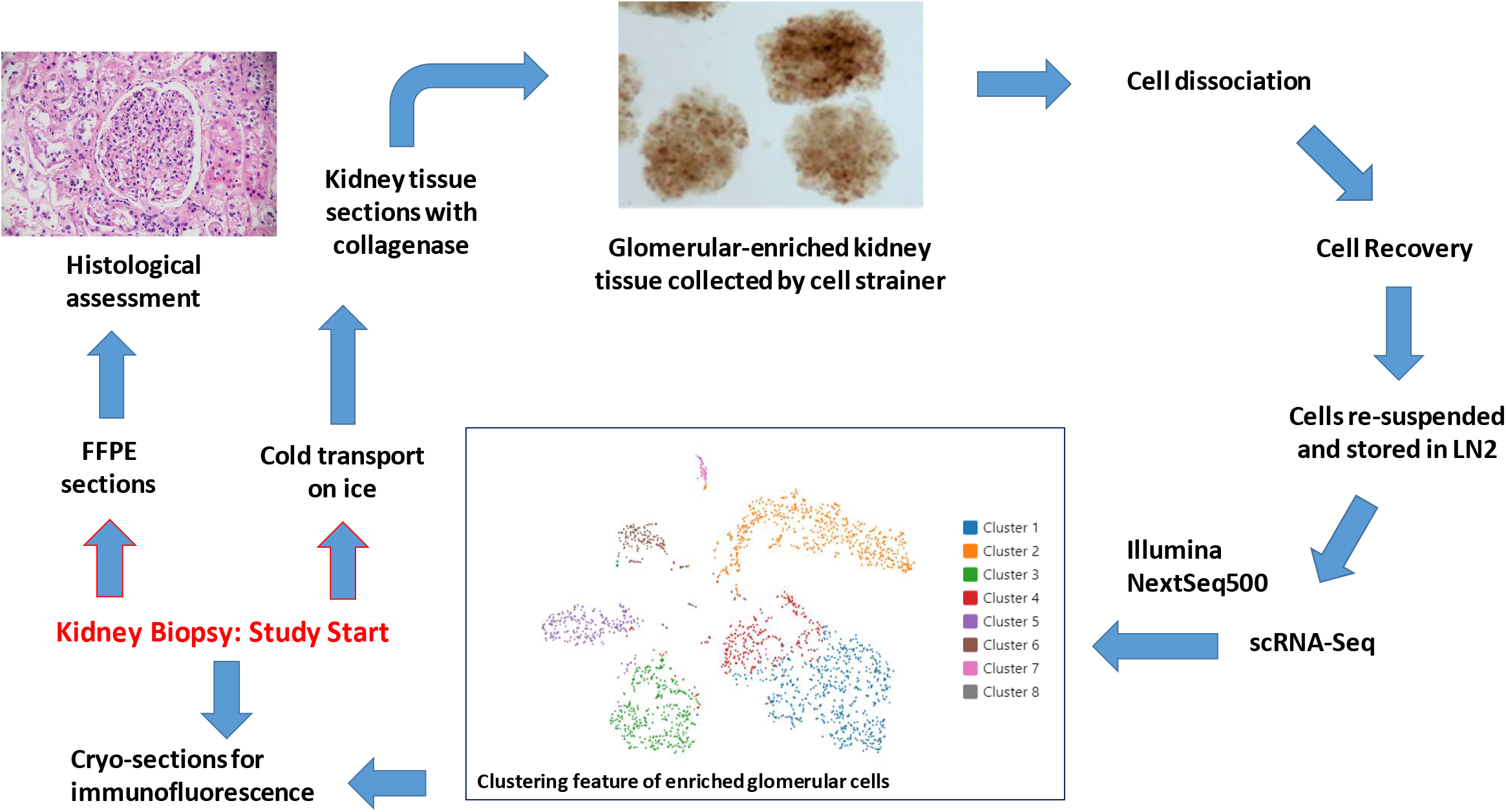
Study design, representative data, and technical outline Note: Photos are actual images; the analytical data shown is from a single representative participant

**Supplementary Figure 2.**
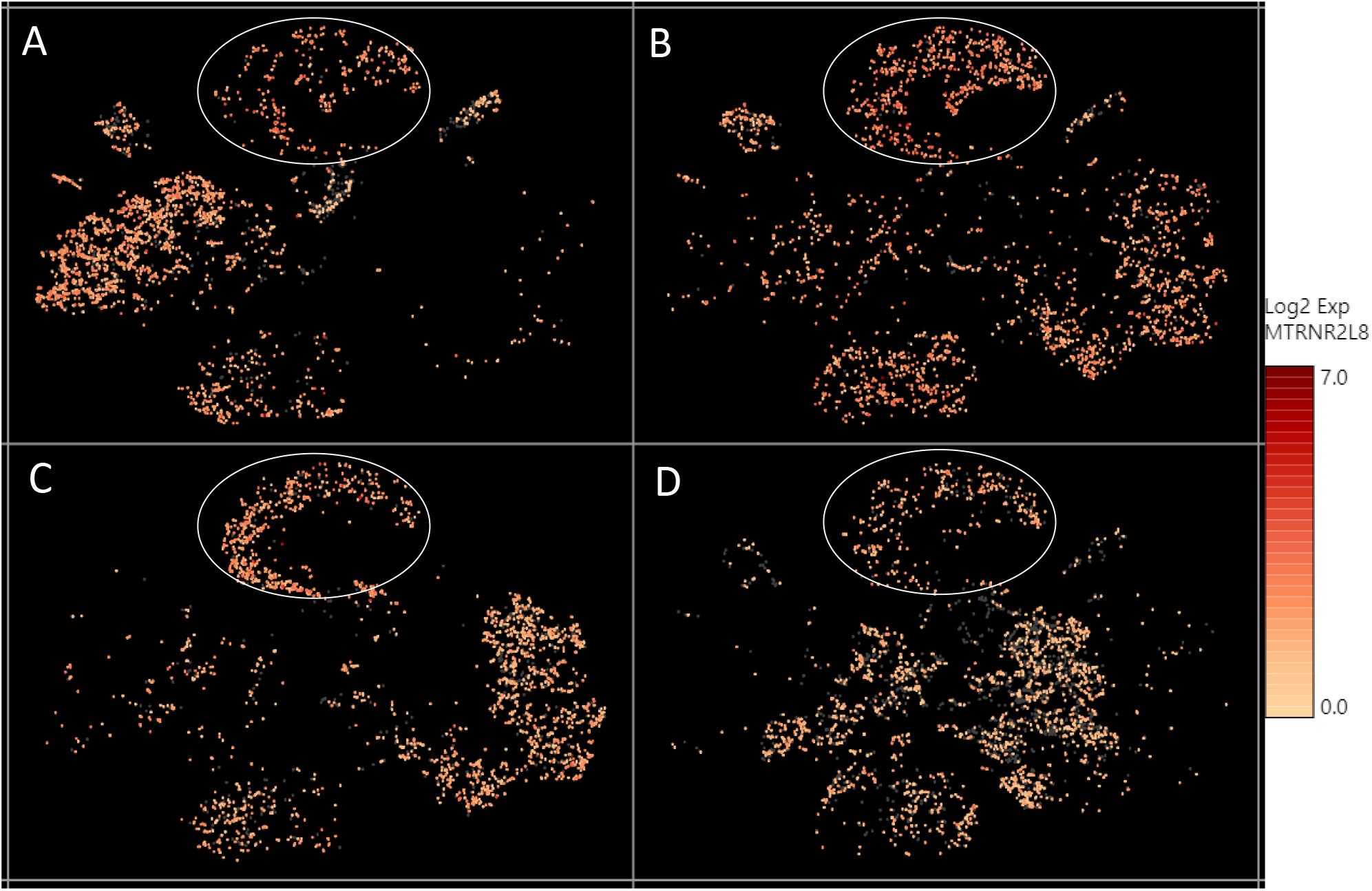
The relative expression level of a mitochondrial encoded transcript *MTRNR2L8* across different cell clusters. *MTRNR2L8* levels were higher in the “undetermined” cells (circled), compared to other cell types

**Supplementary Figure 3.**
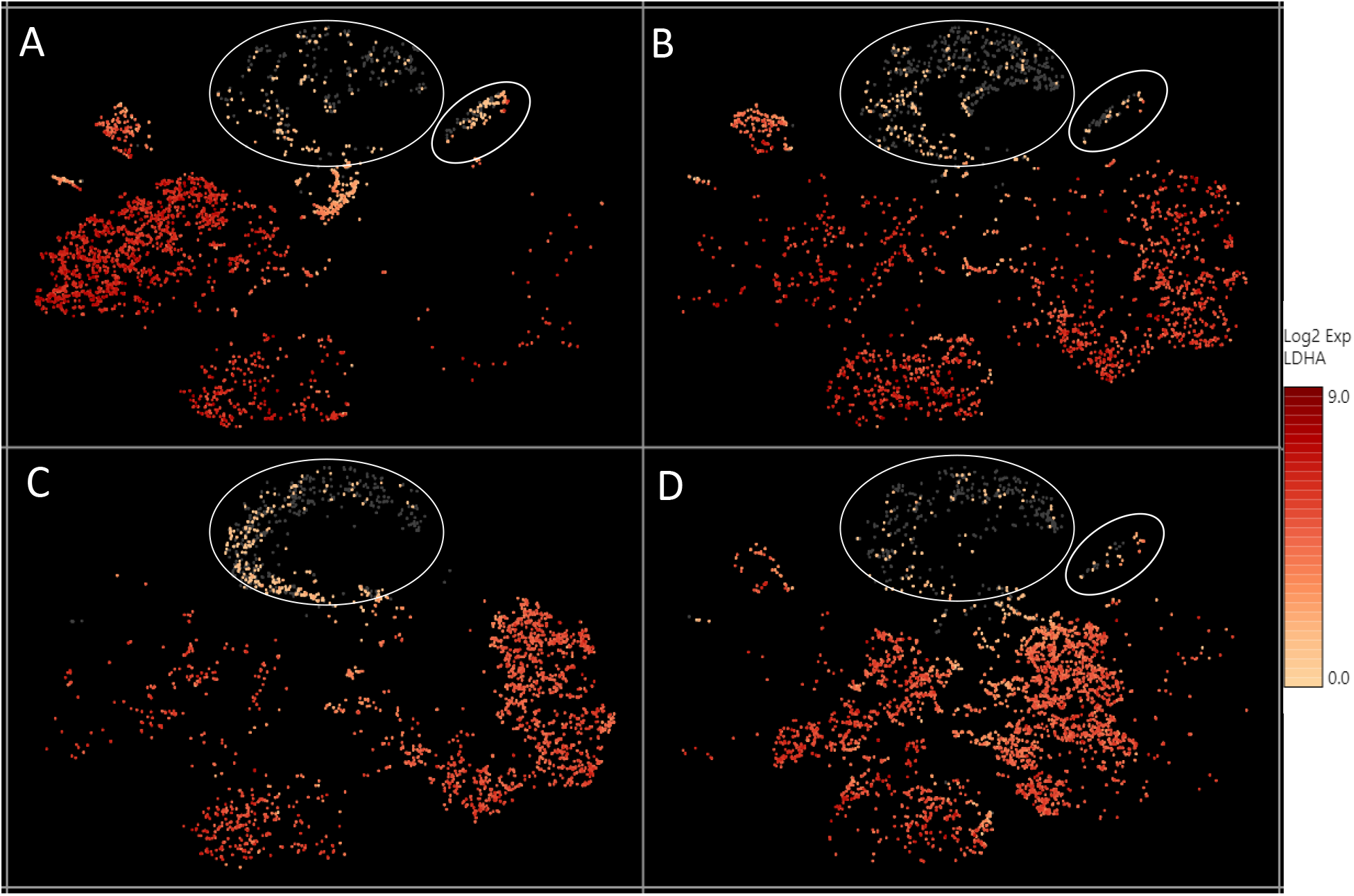
The relative expression level of the *LDHA* transcript across different cell clusters. *LDHA* levels were lower in the “undetermined” and podocyte cell clusters (circled), compared to other cell types

**Supplementary Figure 4.**
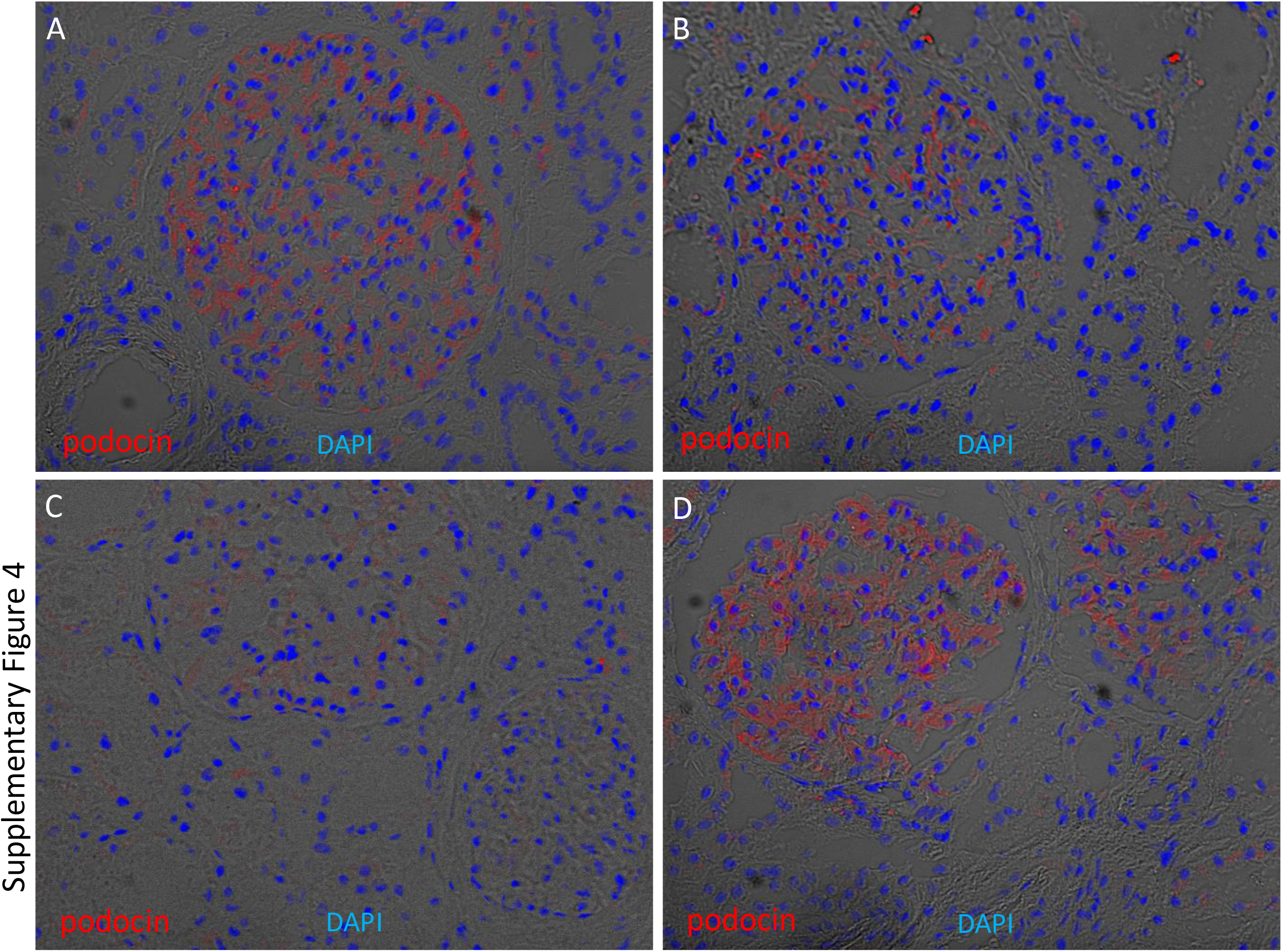
Glomerular enrichment of podocin on kidney cryosections. Immunofluorescence of podocin was performed on four kidney cryosections from European American patients without clinical kidney disease. Cryosections were stained for podocin (red) and counterstained with 4’,6-diamidino-2-phenylindole (DAPI) (blue) overlapped with bright field. Podocin signals are enriched in the glomeruli, with weaker expression in Patient C.

**Supplementary Figure 5.**
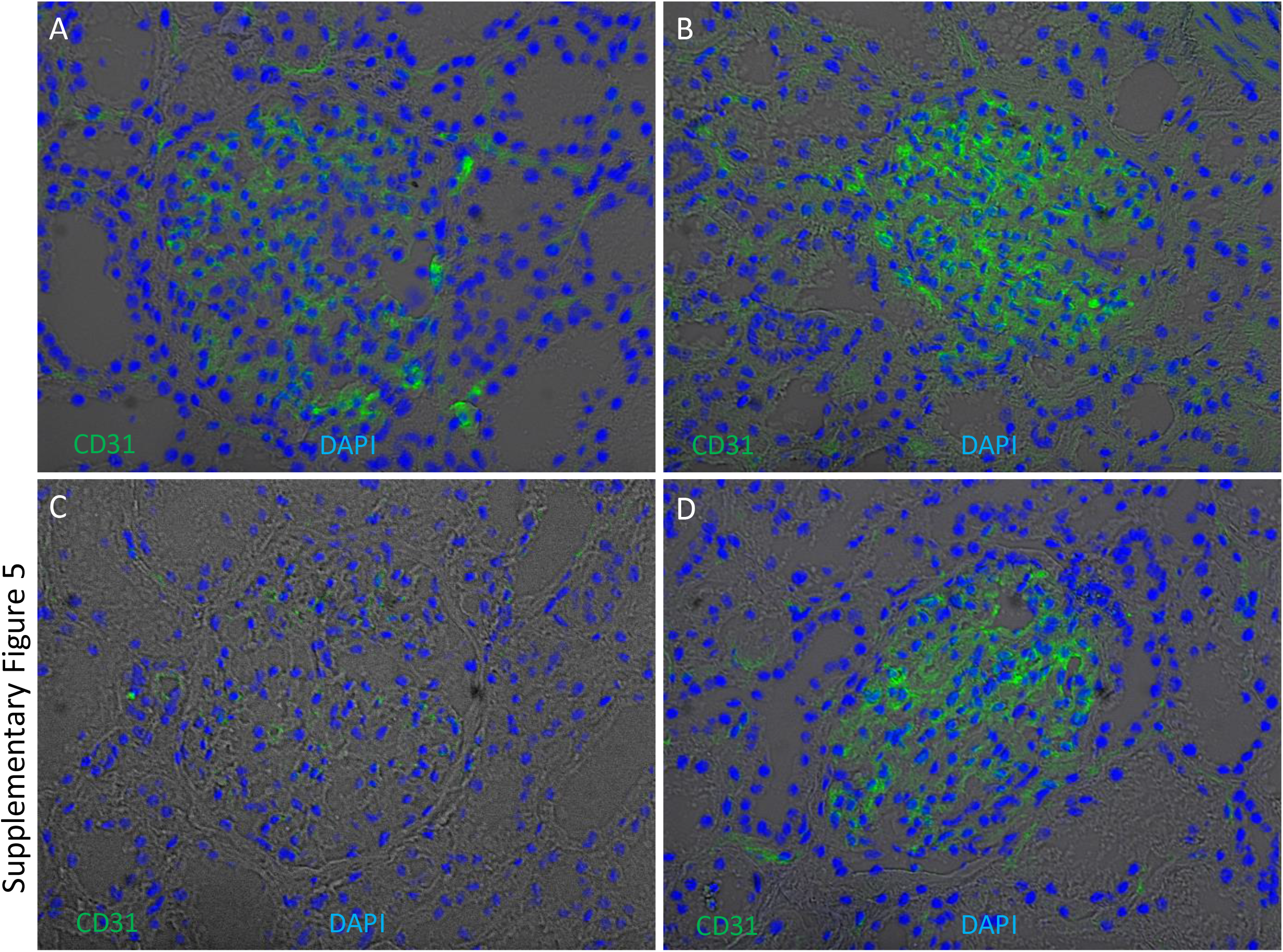
Glomerular enrichment of CD31 (PECAM1) on kidney cryosections. Immunofluorescence of CD31 was performed on four kidney cryosections from European American patients without clinical kidney disease. Cryosections were stained for CD31 (green) and counterstained with 4’,6-diamidino-2-phenylindole (DAPI) (blue) overlapped with bright field. CD31 signals are enriched in glomeruli, with weaker expression in Patient C.

**Supplementary Figure 6.**
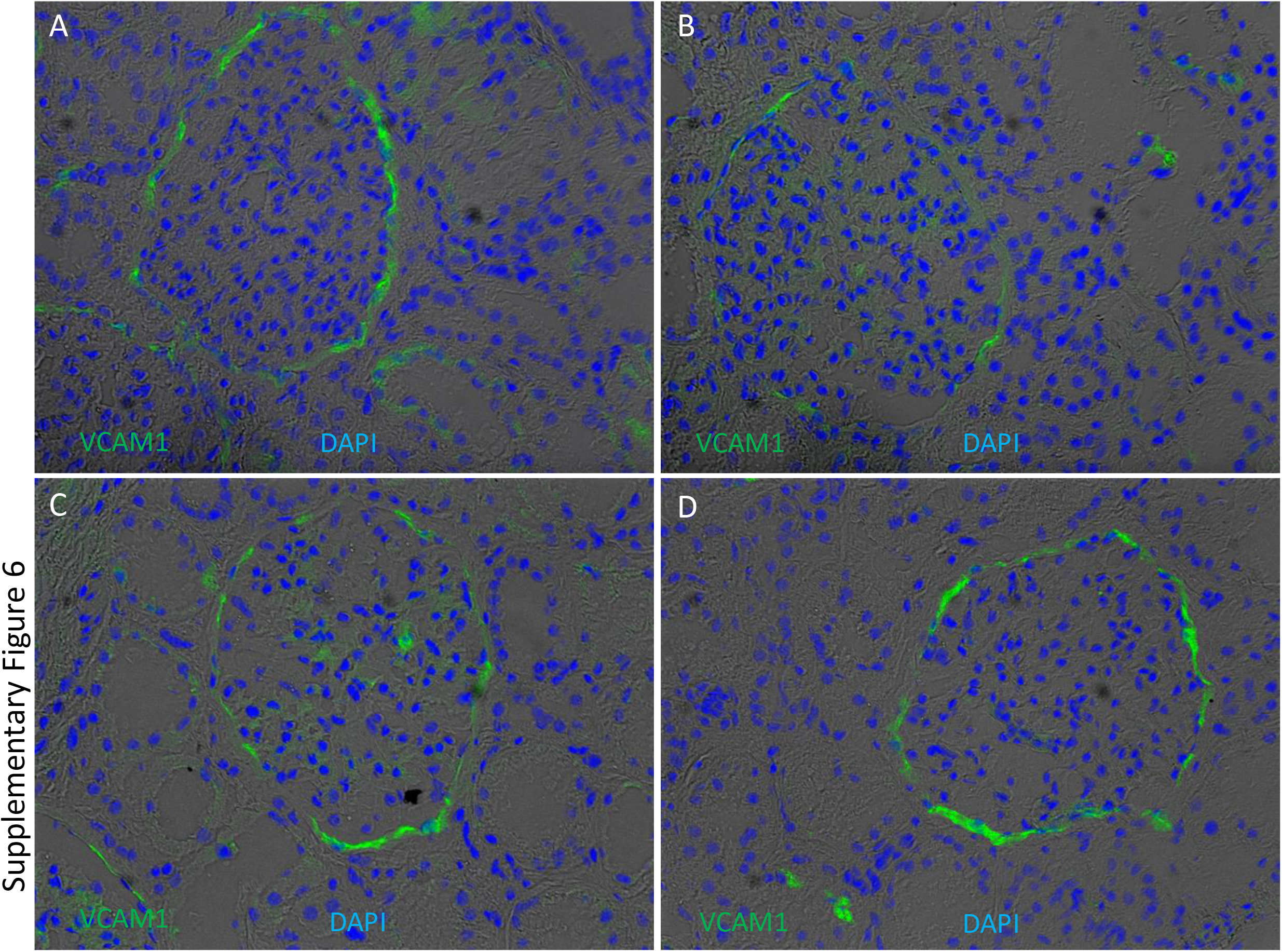
Presence of VCAM1 on kidney cryo-sections. Immunofluorescence of VCAM1 was performed on four kidney cryosections from European American patients without clinical kidney disease. Cryosections were stained for VCAM1 (green) and counterstained with 4’,6-diamidino-2-phenylindole (DAPI) (blue) overlapped with bright field. VCAM1 appeared to enriched in cells lining Bowmen capsule and peritubular endothelial cells.

